# Strain and serovar variants of *Salmonella enterica* exhibit diverse tolerance to food chain-related stress

**DOI:** 10.1101/2022.10.11.511718

**Authors:** Hannah V. Pye, Gaëtan Thilliez, Luke Acton, Rafał Kolenda, Haider Al-Khanaq, Stephen Grove, Robert A. Kingsley

## Abstract

Non-Typhoidal *Salmonella* (NTS) continues to be a leading cause of foodborne illness worldwide. Food manufacturers implement hurdle technology by combining more than one approach to control food safety and quality, including preservatives such as organic acids, refrigeration, and heating. We assessed the variation in survival in stresses of genotypically diverse isolates of *Salmonella enterica* to identify genotypes with potential elevated risk to sub-optimal processing or cooking. Sub-lethal heat treatment, survival in desiccated conditions and growth in the presence of NaCl or organic acids were investigated. *S*. Gallinarum strain 287/91 was most sensitive to all stress conditions. While none of the strains replicated in a food matrix at 4°C, *S*. Infantis strain S1326/28 retained the greatest viability, and six strains exhibited a significantly reduced viability. A *S*. Kedougou strain exhibited the greatest resistance to incubation at 60°C in a food matrix that was significantly greater than *S*. Typhimurium U288, S Heidelberg, *S*. Kentucky, *S*. Schwarzengrund and *S*. Gallinarum strains. Two isolates of monophasic *S*. Typhimurium, S04698-09 and B54 Col9 exhibited the greatest tolerance to desiccation that was significantly more than for the *S*. Kentucky and *S*. Typhimurium U288 strains. In general, the presence of 12mM acetic acid or 14mM citric acid resulted in a similar pattern of decreased growth in broth, but this was not observed for *S*. Enteritidis, and *S*. Typhimurium strains ST4/74 and U288 S01960-05. Acetic acid had a moderately greater effect on growth despite the lower concentration tested. A similar pattern of decreased growth was observed in the presence of 6% NaCl, with the notable exception that *S*. Typhimurium strain U288 S01960-05 exhibited enhanced growth in elevated NaCl concentrations. An understanding of the molecular basis of phenotypic variation in response to stress has the potential to improve process validation during food challenge tests, improve processing, and result in more reliable risk assessments in the food industry.

## 1. Introduction

Non-Typhoidal *Salmonella* (NTS) is a leading cause of foodborne disease worldwide. In the EU, ~88,000 cases of human salmonellosis were reported in 2019 and 17.9% of all foodborne outbreaks were attributable to *Salmonella* (ECDPC, 2021). The food products commonly associated with foodborne salmonellosis outbreaks were eggs, egg products, bakery products, pork and mixed food products (ECDPC, 2021). Salmonellosis typically presents as a self-limiting gastroenteritis with diarrhoea, nausea, abdominal pain and fever, usually occurring between 6 and 48 hours after ingestion of food and drink contaminated with *Salmonella* (Crum-Cianflone, 2008). The minimum infective dose can be between 10^5^ and 10^10^ cells, depending on serovar, associated food type and individual (McCullough and Eisele, 1951a; McCullough and Eisele, 1951b; McCullough and Eisele, 1951c; Eisele and McCullough, 1951). In severe cases, individuals may be hospitalised if bloody diarrhoea or dehydration occurs, sometimes resulting in death in immunocompromised patients if not treated effectively (Hardy, 2004).

Ancestors of the genus *Salmonella* gained virulence factors required to invade epithelial cells and trigger an immune response associated with diarrhoeagenesis, shortly after divergence from the common ancestor with *Escherichia coli* (Baumler, 1997). Consequently, virtually all serovars can cause infection in people. Over 2600 serovars are defined for the genus *Salmonella* but over half of human infections in the United Kingdom in 2019 were caused by strains of *S*. Enteritidis or *S*. Typhimurium, and nearly two thirds were caused by strains of just ten serovars (UKHSA, 2021). The apparent difference in risk to food safety of serovars of the genus *Salmonella* is likely due to a combination of factors, including the opportunity to enter the food chain and the ability to tolerate stresses to survive or replicate in the food chain. An important factor in the opportunity to enter the food chain is the host range of the *Salmonella* variant. Most non-typhoidal *Salmonella* are considered broad host range, since they can circulate and cause disease in multiple hosts. Nonetheless, considerable host adaptation of *Salmonella* serovars is evident from the unique complement with little overlap of serovars isolated from different livestock species (APHA, 2021). *S*. Typhimurium, including its monophasic variant, is unusual among *Salmonella* serovars in its wide distribution in many livestock species and indeed wild mammals and birds (Branchu *et al.*, 2018; Rabsch *et al.*, 2002). Extreme cases of host restriction may also play a role in the absence of some serovars from human infections. For example, *S*. Gallinarum only circulates in poultry and is virulent in other species for reasons that are not fully understood. Other highly host adapted serovars such as *S*. Choleraesuis and *S*. Dublin that are adapted to circulation in pig and bovine hosts where they cause a more severe disseminated disease, can also cause disease in people, but these infections are relatively rare (Kingsley and Bäumler, 2000; Stevens and Kingsley, 2021).

A second factor that may affect risk to food safety is the ability of serovars to survive or replicate in the food chain. The ability of *Salmonella* to survive and replicate during stress enables *Salmonella* to persist in food and food processing environments. Stress may be defined as any physical, nutritional or chemical process which results in sub-lethally injured bacteria (Wesche *et al.*, 2009). Stresses include high and low temperature, desiccation, and chemical preservatives such as organic acids and salt. Desiccation is used as a food preservation technique to extend the shelf-life of products and these products typically have a low-water activity (a_w_ < 0.85), which is the ratio of water vapour pressure in a food product compared to the water vapour pressure of pure water at a specified temperature (Finn *et al.*, 2013; Labuza, 1980). Salt is also commonly used as a preservative and works by preventing bacterial growth through lowering the water activity of food. *Salmonella* survives osmotic stress by increasing the availability of intracellular solutes, which are then replaced by proline, trehalose and glycine betaine to initiate growth (Koo and Booth, 1994). Thermal treatment, using water, steam, or thermal radiation, is one of the most effective methods of eliminating foodborne pathogens. During pasteurisation, a heat treatment of between 65°C and 95°C is applied to the food product to inactivate vegetative pathogens (Silva and Gibbs, 2009). Exposure to high temperatures (> 60°C) causes cytoplasmic ions to leak out of the cell, and irreversible damage to the cell membrane resulting in cell death (Ebrahimi *et al.*, 2018). Low temperatures inhibit microbial growth by reducing membrane fluidity, and disrupting cellular processes, such as mRNA transcription and translation and triggers protein degradation (Phadtare, 2004). Weak organic acids such as lactic, acetic and citric acid are used as preservatives in food manufacturing and can be defined as organic compounds that retain acidic properties (Sauer *et al.*, 2008). They are normally bacteriostatic rather than bactericidal, by diffusing through the cytoplasm and dissociating into protons and anions on exposure to the neutral pH inside the cell. Acidification constrains cellular functions due to inhibition of acid-sensitive enzymes such as those involved in glycolysis (Lambert and Stratford, 1999).

Modern food manufacturing practices rarely rely on one form of preservation to inhibit microbial growth in food products, rather, a combination of multiple techniques are commonly used to provide robust protection against food spoilage and pathogenic microorganisms (Leistner, 2000). However, consumer demand for minimally processed food is increasing, and therefore understanding how pathogens behave under stress will aid in developing processing techniques which satisfy consumer interests for healthier products, without compromising food safety. To assess the effectiveness of preservation and exclusion of pathogens, typically, multiple *Salmonella* strains are used together in a strain cocktail to assess survival during food challenge tests, but certain strains possess an increased resistance to stress, thus posing a greater risk to food safety. Understanding strain variation will improve modelling of consumer risk of contamination of *Salmonella* in food products (Whiting and Golden, 2002) and improve selection of target strains for process validations, improved processing, and more reliable risk assessments. This study aims to evaluate the phenotypic variability amongst *Salmonella* strains in response to food chain related stress, and is the first study, to our knowledge, to evaluate multiple stresses on the same subset of strains, identifying key serovars with an increased resistance to food chain related stress.

## 2. Methods

### 2.1 Bacterial Strains and Culture

Fourteen *S. enterica* strains primarily isolated from food producing animals and food products were used in the present study. These included five strains of serovar Typhimurium, two strains of serovar Newport and a single representative strain of Kedougou, Infantis, Heidelberg, Enteritidis, Kentucky, Gallinarum and Schwarzengrund. Stock cultures of each strain were stored in individual Cryovials (Corning) at −80°C in 50% Glycerol. Working cultures were prepared by incubation in 5mL Luria-Bertani (LB) broth at 37°C with shaking at 200rpm for 18 hours. Serovars to be included in this study were selected due to their invasiveness, their ability to cause disease in humans and food production animals, or because they were isolated from food or food production environment (Table 1). A range of serovars were also chosen to be included from across the phylogeny, to incorporate as much genetic diversity as possible.

**Table 1.** *S. enterica* strains used in this study.

| serovar | sequence Type | phage type* | strain ID | source | sequence accession |
| --- | --- | --- | --- | --- | --- |
| Typhimurium | ST19 | ND | ST4/74 | Cattle | SAMN30870364 |
| Typhimurium | ST568 | DT56 | S07676-03 | Avian | SAMN30870365 |
| Typhimurium | ST19 | U288 | S01960-05 | Pig | ERS015663 |
| Typhimurium | ST34 | DT193 | S04698-09 | Cattle | ERS023534 |
| Typhimurium | ST34 | ND | B54Col9 | Chicken | SAMN30870366 |
| Kedougou | ST1543 | NA | B37Col19 | Cattle | SAMN30870367 |
| Infantis | ST32 | NA | S1326/28 | Chicken | SAMN30870368 |
| Heidelberg | ST15 | NA | SL476 | Turkey | SAMN30870369 |
| Enteritidis | ND | PT4 | P125109 | Human | SAMN30870370 |
| Schwarzengrund | ST322 | NA | SL480 | Human | SAMN30870371 |
| Gallinarum | ND | NA | 287/91 | Chicken | SAMN30870372 |
| Newport | ST45 | NA | SL254 | Human | SAMN30870373 |
| Newport | ST375 | NA | SGSC4157 | Unknown | SAMN30870374 |
| Kentucky | ST152 | NA | SL479 | Human | SAMN30870375 |

\* ND – not determined, NA – not applicable

### 2.2 Preparation and storage of vegetarian food product

Individual packets of a wheat and pea protein-based vegetarian food product (each pack totalling 210g) were subjected to an in-pack irradiation treatment and were supplied by Nestlé for use throughout the study. The 210g packets of food product remained at 4°C until opened, and once opened, each pack was divided into 13g portions in individual sterile plastic bags and then frozen at −20°C. Individual portions of food product were thawed at room temperature prior to use. The product was homogenised by hand and used in stress response assays.

### 2.3 Whole Genome Sequencing and Phylogenetic analysis

*Salmonella* strains were cultured overnight at 37°C in LB broth and DNA was extracted using a Maxwell RSC 48 instrument (Promega) and associated Maxwell RSC cultured cells DNA kit (Promega). DNA concentration was quantified using the Qubit 3.0 fluorometer for dsDNA broad range assay kit (Thermo Fisher Scientific). Genomic libraries were prepared using the Nextera XT index kit (Illumina) and whole genome sequencing was performed using a NextSeq500 (Illumina). Data was uploaded to IRIDA (version 19.09.2) (Matthews *et al.*, 2018) and quality checked using FastQC (version 0.11.9) (Andrews, 2010). Antigenic formula was predicted using SeqSero2 (version 1.2.1) using raw forward and reverse short read sequences to identify serotype (Zhang *et al.*, 2019). SNIPPY (version 4.3.6) (Seemann, 2015) was used to identify single nucleotide polymorphisms (SNPs) between *Salmonella* strains used in this study and the *S. bongori* N268-08 reference. A maximum likelihood phylogenetic tree was constructed from the core alignment from the SNIPPY output using RaxML (Stamatakis, 2014) and plotted using ggtree (Yu *et al.*, 2017) in R (version 4.1) (Team, 2021). FastANI (version 1.3) was used to determine the average nucleotide identity (ANI) between strains, using the many-to many method, where multiple query and reference genomes were used (Jain *et al.*, 2018). The ANI between strains was plotted as a matrix with heatmap3 (version 1.1.9) (Zhao *et al.*, 2021) in R (version 4.1) (Team, 2021), with the parameter (symm = T) to maintain symmetry.

### 2.4 Heat Inactivation of *Salmonella enterica*

*Salmonella enterica* strains were grown for 18 hours in 5mL LB broth at 37°C and then pelleted by centrifugation at 13,300rpm for 4 minutes. The supernatant was removed, and the resulting bacterial pellet resuspended in an equal volume of phosphate buffered saline (PBS). Strains were diluted to approximately 5×10^8^ CFU/mL with PBS and stored at 2-4°C for 1 hour. Individual thin-walled water-tight aluminium thermal cells (supplied by Nestlé) were filled with 750mg of thawed vegetarian food product and inoculated with 5μL of each *Salmonella* strain at approximately 5×10^8^ CFU/mL. Inoculated thermal cells were refrigerated at 2-4°C for 1 hour to allow the cells to equilibrate within the food matrix. Thermal cells were heated in a waterbath at 60°C for 30 seconds once the temperature at the centre of the thermal cells reached the required temperature (~45 seconds). The temperature of food matrix was monitored throughout the experiment by attaching at least one thermal cell containing inoculated product to a thermocouple (type K) and a TC-08 Temperature Data Logger (Pico Technology). Once heated, thermal cells were plunged in an iced water bath to rapidly cool. For a control, thermal cells were inoculated in the same way but remained at room temperature. The contents of each thermal cell were transferred to a 15mL centrifuge tube (Corning) and vortexed with 7.5mL PBS. A 300μL aliquot of each centrifuge tube was deposited into the first column of a 96-well CytoOne plate (Starlab) and serially diluted (1:10) with PBS. A 5μL aliquot of each dilution was spotted onto a square 100mm plate (Thermo Scientific) containing 50mL LB agar (in triplicate). Surviving colonies were enumerated after overnight incubation at 30°C and the log ratio survival determined. Five independent experiments were conducted for each strain at 60°C.

### 2.5 Survival at Refrigerated Temperatures

Strains were grown to stationary phase in 5mL LB broth for 18 hours at 37°C and then pelleted using centrifugation at 13,300rpm for 4 minutes. The supernatant was discarded, and the pellet resuspended in an equal volume of PBS. Cultures were adjusted to approximately 2.5×10^9^ CFU/mL and refrigerated at 2-4°C for 1 hour. Wells of a 24-well CytoOne plate (Starlab) were filled with 750mg of vegetarian food product and inoculated with 50μL of each strain at approximately 2.5×10^9^ CFU/mL. Plates were sealed using laboratory sealing film (parafilm) and maintained at 4°C for 5 weeks. An initial assessment of viable counts that could be recovered for each strain was determined by immediately transferring the well contents into 5mL LB broth and viable counts determined by plating serial dilutions (1:10) with PBS. For CFU/mL counts, 5μL of each dilution was spot plated (in triplicate) onto LB agar and incubated at 30°C overnight. After 5 weeks, the contents of the wells from the experimental plates were transferred to LB broth, and viable counts enumerated by spot plating 5μL of each dilution (in triplicate) onto LB agar and incubating at 30°C overnight. Four independent experiments were conducted for each strain from two different overnight cultures.

### 2.6 Desiccation

Strains were cultured for 18 hours in 5mL LB broth at 37°C with shaking at 200rpm. Overnight cultures were centrifuged at 13,300rpm for 4 minutes and the supernatant discarded. The pellet was resuspended in an equal volume of PBS and adjusted to approximately 5×10^8^ CFU/mL. The first column of a 96-well plate (CytoOne) was filled with 50μL of each strain at 5×10^8^ CFU/mL. Plates were left to desiccate in a safety cabinet for 24-hours for complete evaporation and the temperature and relative humidity was measured using a thermohygrometer. Control wells containing 50μL of each strain at approximately 5×10^8^ CFU/mL were mixed with 150μL PBS per well, and 5μL of serial dilutions spot plated, in triplicate, onto LB agar plates and incubated at 30°C for 18 hours. The following day, surviving colonies were enumerated. After 24-hours, desiccated wells were rehydrated with 200μL PBS and serially diluted (1:10) with PBS. A 5μL aliquot of each dilution was spotted onto an LB agar plate (in triplicate) and incubated overnight at 30°C. Surviving colonies were counted, and the log ratio survival calculated. Three independent experiments were conducted for each strain.

### 2.7 Growth in the presence of organic acids

*Salmonella enterica* strains were cultured for 18 hours in 5mL LB broth at 37°C with shaking at 200rpm. A 14mM citric acid (Thermo Scientific) solution and a 12mM acetic acid (Scientific Laboratory Supplies) solution were prepared in LB broth and filter sterilised using a 0.2μM minisart PES syringe filter (Sartorius). A 1mL aliquot of overnight culture of each strain was mixed with 4mL LB broth supplemented with either 14mM citric acid or 12mM acetic acid at pH 5.8 (adjusted using the appropriate acid) for 30 minutes, to initiate the acid shock tolerance response. This methodology was used to improve consistency of results between biological replicates and to condition cells to survive in a low pH, which would be most similar to the pH used in food products to prevent microbial growth. 5mL aliquots of organic acid-supplemented LB broth solutions were inoculated with 5μL of each test strain that had been pre-adapted to pH 5.8 at a concentration of approximately 5×10^8^ CFU/mL to give a final concentration of approximately 5×10^5^ CFU/mL. A 200μL aliquot of inoculated organic acid solution for each strain was transferred to a 96-well U-Bottom plate (Greiner), in triplicate wells. LB broth (positive) controls for each strain and non-inoculated (negative) controls were both included. Growth was measured at OD600nm using a FLUOstar Omega plate reader (BMG Labtech) for 22 hours, with measurements taken every 5 minutes at 37°C with pre-measurement shaking. Three independent experiments were conducted for each strain. Area under the curve calculations were conducted in GraphPad Prism (version 8.0.2) from the average of each biological replicate. An R package (version 4.1), called Growthcurver, was used to determine the maximum OD600nm, and generation time (t_gen) for all 14 strains included in the study.

### 2.8 Growth in the presence of salt

*Salmonella enterica* strains were grown to stationary phase overnight at 37°C in 5mL LB broth. Cultures were diluted to approximately 5×10^8^ CFU/mL with LB broth. 5mL LB broth containing 6% NaCl was aliquoted and inoculated with 5μL of each strain at approximately 5×10^8^ CFU/mL. Each well of a 96-well U-Bottom plate was filled with 200μL of inoculated salt solution for each strain (in triplicate). Non-inoculated LB broth controls (with and without NaCl), and inoculated LB broth controls were included. Growth was measured at OD600nm using a FLUOstar Omega plate reader (BMG Labtech) for 22 hours, with measurements taken every 5 minutes at 37°C with pre-measurement shaking. Three independent experiments were conducted for each strain. Area under the curve calculations were conducted in GraphPad Prism (version 8.0.2) from the average of each biological replicate. An R package (version 4.1), called Growthcurver, was used to determine the maximum OD600nm, and generation time (t_gen) for all 14 strains included in the study.

### 2.9 Statistical Analysis

A one-way ANOVA, with Fisher’s least significant difference (LSD) test, was conducted in Graphpad prism (version 8.0.2) on the log ratio survival data of each replicate mean compared to the mean of the most resistant strain for each stress condition. A two-way ANOVA with multiple comparisons and Fisher’s Least significant difference test was conducted on the area under the curve (AUC) analysis in Graphpad Prism (version 8.0.2) comparing the mean of each stress compared to the mean of the most resistant strain.

## 3. Results

### 3.1 Phylogenetic relationship of *S. enterica* strains

*S. enterica* strains used during the study were selected to represent a genetically diverse collection of serotypes from *Salmonella enterica* subspecies I and strains with diverse host range and association with food or livestock. The diversity and phylogenetic relationship of *S. enterica* strains used throughout this study was determined based on nucleotide sequence variation in the shared genome. A maximum likelihood tree indicated a star topology with most of the strains of distinct serotypes present on long branches with common ancestors close to root, defined by *S*. *bongori* strain N268-08 as an outgroup (Figure 1A), consistent from previous reports (Branchu *et al.*, 2018). *S.* Enteritidis strain P125109 and *S.* Gallinarum strain 287/91 were present on short terminal branches indicating a relatively recent common ancestor and similar core genome sequences (Thomson *et al.*, 2008). Five strains of *S*. Typhimurium clustered closely together and were present on very short terminal branches as reported previously (Bawn et al., 2020; Branchu *et al.*, 2018). Average sequence identity of strains in distinct phylogroups that contained distinct serotypes ranged from 98.13% for *S*. Schwarzengrund and *S*. Newport strain SGSC4157 to 99.7% for *S.* Enteritidis strain P125109 and *S.* Gallinarum strain 287/91, with a mean of 98.89%. (Figure 1B). Five strains of *S*. Typhimurium used in this study had an ANI > 99.8%.

**Figure 1.**
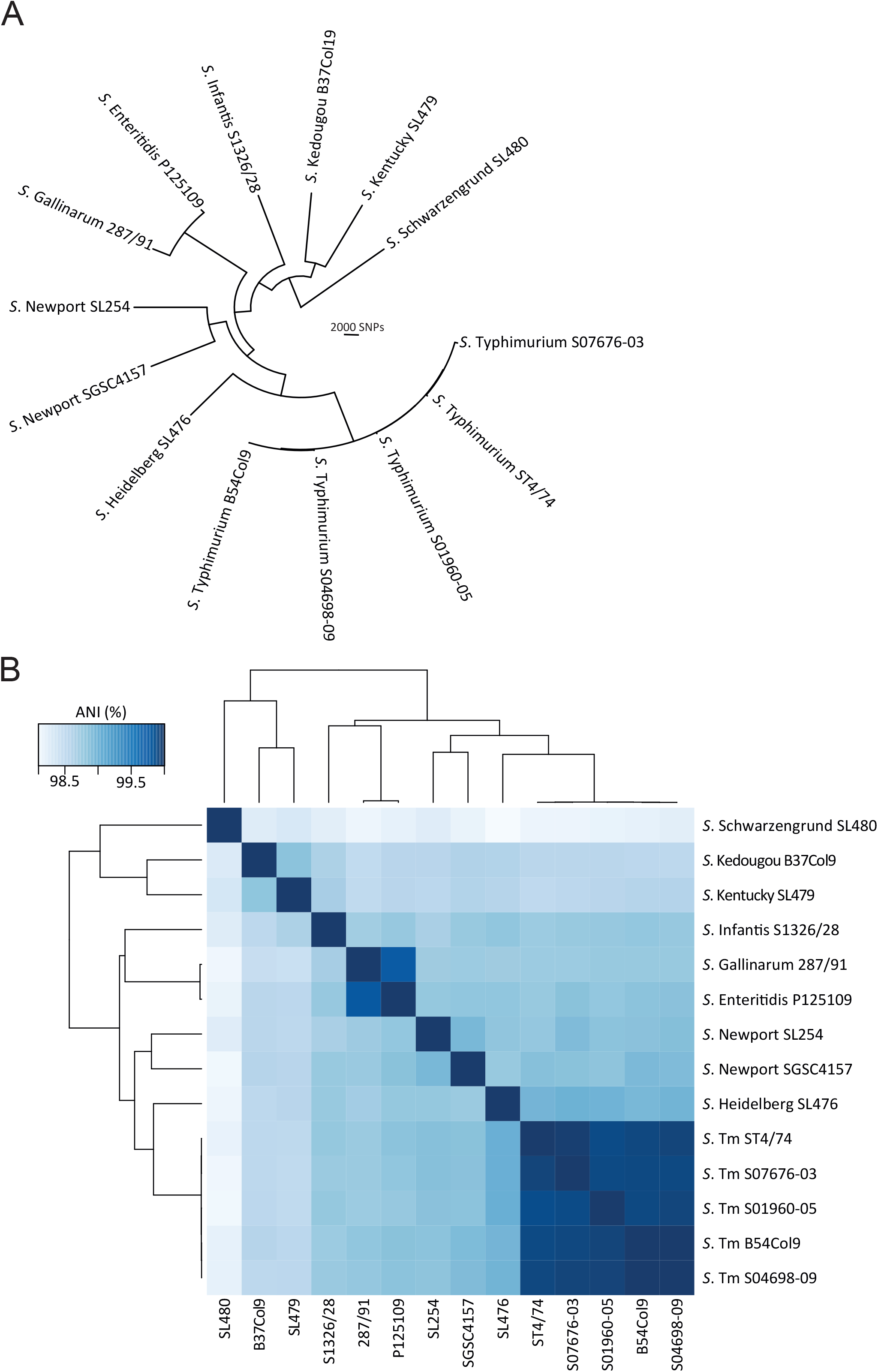
Phylogenetic relationship of *S. enterica* strains used in this study. (A) Maximum likelihood phylogenetic tree of *Salmonella enterica* subspecies I strains used in this study based on variation in the core genome. The tree is rooted using *S*. *bongori* as an outgroup (not shown). (B) Average nucleotide identity (ANI) calculated for pairs of genomes using FastANI values represented as a heatmap. The percentage identity is represented by the intensity of colour indicated by the key (inset).

### 3.2 *Salmonella* genotype affects survival in extremes of temperature and during desiccation

*Salmonella* may encounter multiple stresses during processing, storage or suboptimal cooking, including extremes of temperature and desiccation. To investigate the variation in survival for each strain to these stresses we determined the change in viable counts following storage at 4°C for five weeks or heating to 60°C for 30 seconds in a vegetable-based product, or desiccation by air drying.

The variation in the ability of *Salmonella* to survive for five weeks in a vegetarian food matrix was assessed at 4°C to emulate the products shelf-life and typical storage conditions. No increase in viable counts was observed for any of the strains tested, indicating a lack of net replication (Figure 2A). *S*. Infantis strain S1326/28 exhibited the greatest tolerance to storage with a mean reduction in viable counts of 0.04 log. Most strains showed little or no decrease in viable counts but *S.* Kentucky strain SL479, *S*. Typhimurium strain ST4/74, *S*. Heidelberg strain SL476, *S*. Newport strain SGSC4157 and *S*. Gallinarum strain 287/91 were all recovered at significantly lower levels (p<0.05) compared to *S*. Infantis strain S1326/28 (Figure 2A). *S*. Gallinarum strain 287/91 exhibited the greatest reduction of approximately 0.6 log in cell viability.

**Figure 2.**
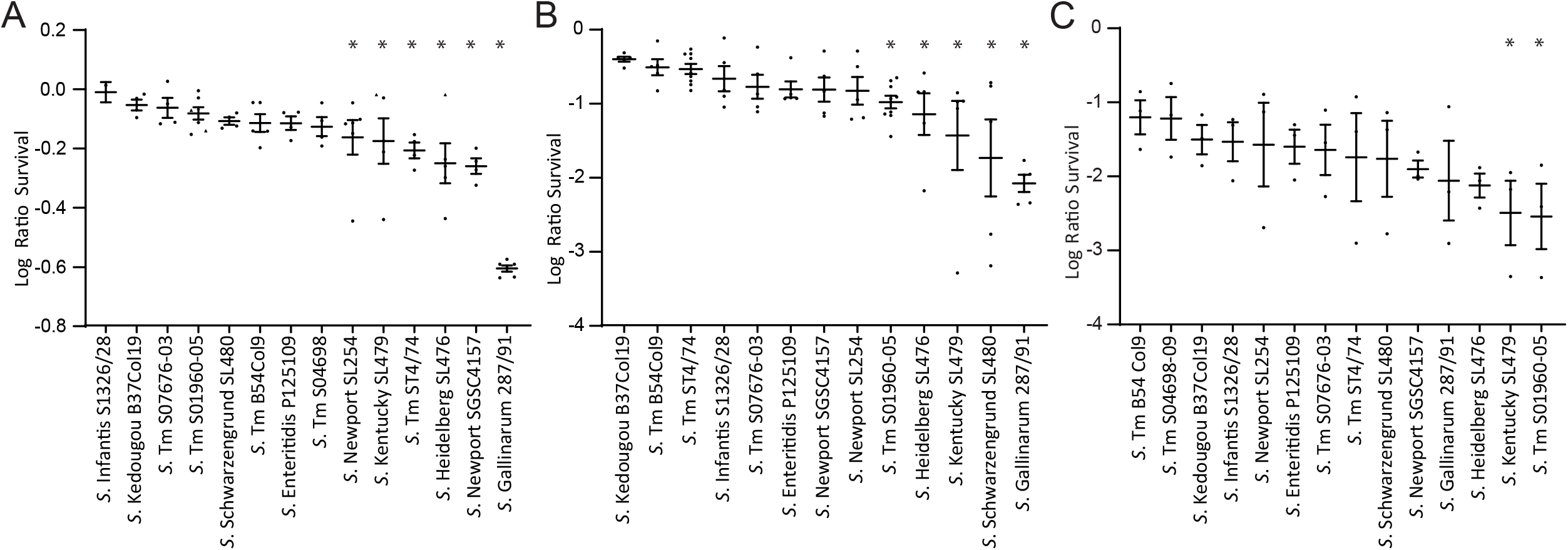
Survival of *Salmonella enterica* strains following cold storage or heat inactivation in food matrix or desiccation in polystyrene wells. (A) Log ratio survival of *Salmonella enterica* strains after storage at 4°C for 5 weeks, in the vegetarian food matrix. Individual points represent each technical replicate, and the symbol represents the biological replicate. (B) Log ratio survival of *Salmonella* strains after heat inactivation at 60°C for 30 seconds in a thermal cell containing 750mg of vegetarian food product. Each Individual point represents a biological replicate. (C) Log ratio survival of *Salmonella* strains after desiccation in a safety cabinet for 24 hours at an average relative humidity of 39% and an average temperature of 21°C. Each biological replicate is plotted as a separate point and represents the mean of 5 technical replicates.

Heat inactivation of *Salmonella* may occur as part of food production or during cooking. To identify suitable conditions that replicate sub-optimal heat treatment or cooking we investigated the inactivation kinetics of *S*. Typhimurium strain ST4/74 in a vegetable-based food matrix. We therefore tested heat inactivation of *Salmonella* strains in a vegetarian food matrix at 60°C for 30 seconds using an aluminium thermal cell (Figure 2B). *S*. Kedougou exhibited the greatest thermal tolerance with this and seven additional strains exhibiting less than a 10-fold decrease in viable counts. Viable counts of five strains, *S*. Typhimurium S01960-05 (U288), *S*. Kentucky strain SL479, *S.* Schwarzengrund strain SL480 and *S*. Heidelberg strain SL476, were each reduced significantly more (p<0.05) than that observed for *S*. Kedougou. *S*. Gallinarum was the most heat sensitive strain with viable counts decreased by over 100-fold.

*Salmonella* can persist in low-moisture food products for extended periods of time, therefore, to assess the variability in desiccation stress tolerance, *Salmonella* strains were subjected to desiccation at an average 39% relative humidity at 21°C. Viable counts for all strains reduced by greater than a factor of 10. Two strains of monophasic *S*. Typhimurium ST34 exhibited the greatest tolerance to desiccation and *S*. Typhimurium strain S01960-05 (U288) and *S.* Kentucky strain SL479 strain had a significantly lower survival (p<0.05) where the reduction in viability exceeded a factor of over 100. (Figure 2C).

### 3.3 *Salmonella* genotype affects growth kinetics in the presence of chemical stress

To assess the variation in growth of each strain in LB broth compared to LB broth supplemented with either NaCl, acetic acid or citric acid, we first identified the concentration at which moderate reduction in growth rate was observed for *S*. Typhimurium strain ST4/74 (supplementary figure 1). A concentration of either 6% NaCl, 12mM acetic acid or 14mM citric acid resulted in a decreased growth of *S*. Typhimurium strain ST4/74 and were used for assessment of all strains by comparing the difference in AUC (Figure 3), the generation time (hours) and maximum OD600nm attained compared with growth in LB broth (Figure 4). *S*. Gallinarum strain 287/91 exhibited the greatest difference in the area under the curve for all three stresses. *S*. Enteritidis strain P125109 and *S*. Typhimurium strain S01960-05 grew better in LB broth containing with 6% NaCl and 14mM citric acid, as observed by a greater area under the curve in the stress condition compared to the control (Figure 3). *S*. Gallinarum strain 287/91 grown in LB broth had the shortest log-phase and the quickest doubling time out of all the trains tested, at ~50 minutes (Figure 4). *S*. Kedougou strain B37 Col19 reached the greatest OD600nm in LB broth (cells pre-adapted to acetic and citric acid) and in LB broth containing 12mM acetic acid. *S.* Enteritidis strain P125109 reached the greatest OD600nm in LB broth containing 14mM citric acid (Figure 4B). *S.* Kentucky strain SL479 exhibited the shortest log-phase and the shortest doubling time in LB broth with 12mM acetic acid, whereas *S*. Newport strain SGSC4157 expressed the shortest log-phase and doubling time (48 minutes) in LB broth with 6% NaCl (Figure 4).

**Figure 3.**
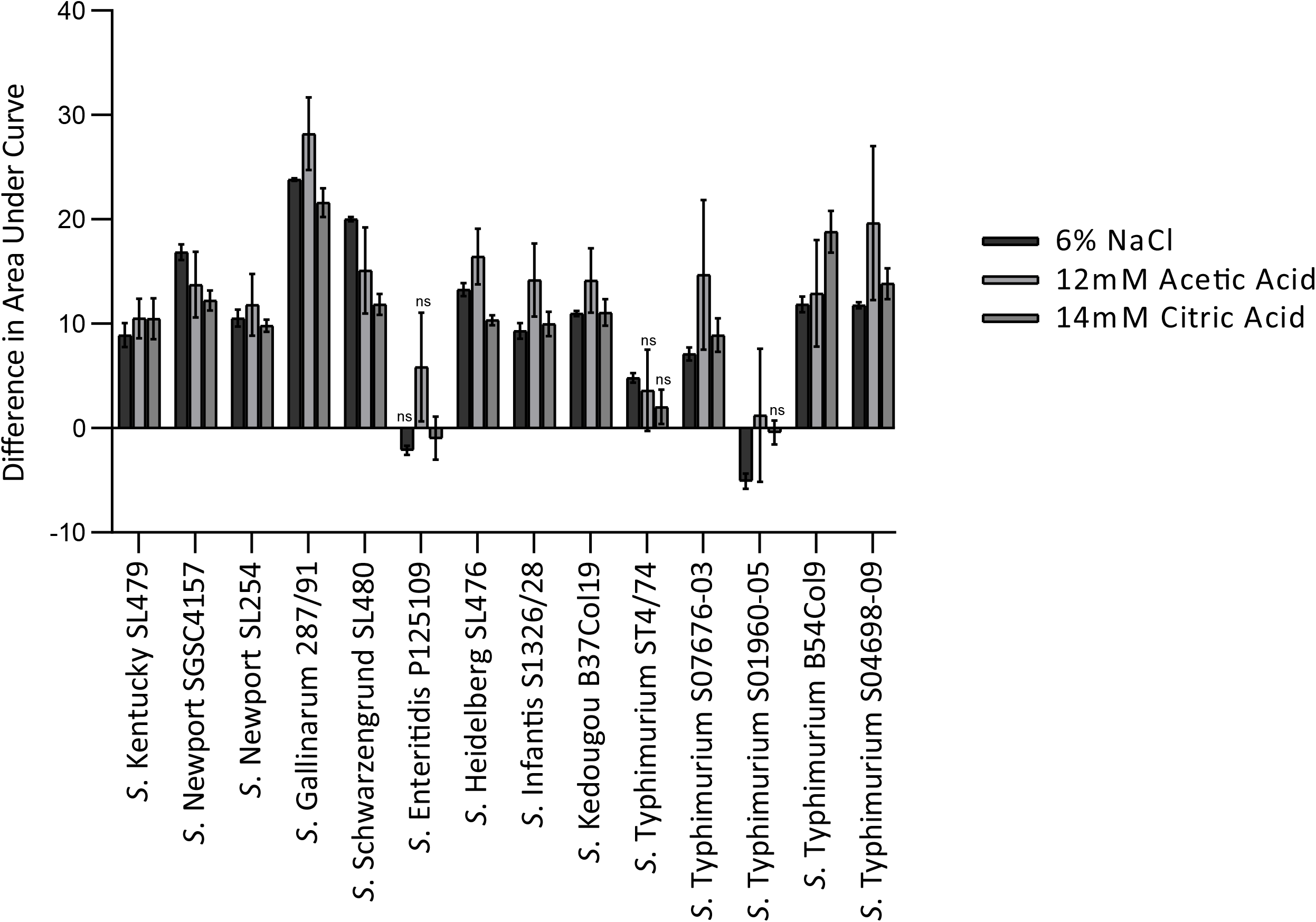
Comparison of growth of *Salmonella enterica* strains in Luria-Burtani broth with or without supplementation with 6% NaCl, 12mM acetic acid or 14mM citric acid. Difference in area under the curve (AUC) for 14 *Salmonella enterica* strains grown in LB broth compared to LB broth supplemented with 6% NaCl, 12mM acetic acid or 14mM citric acid. Bars represent the mean of three biological replicates (±SE). All AUC values were significantly different from the most resistant strain (p < 0.05, two-way ANOVA test) except the ones denoted with ns (not significant).

**Figure 4.**
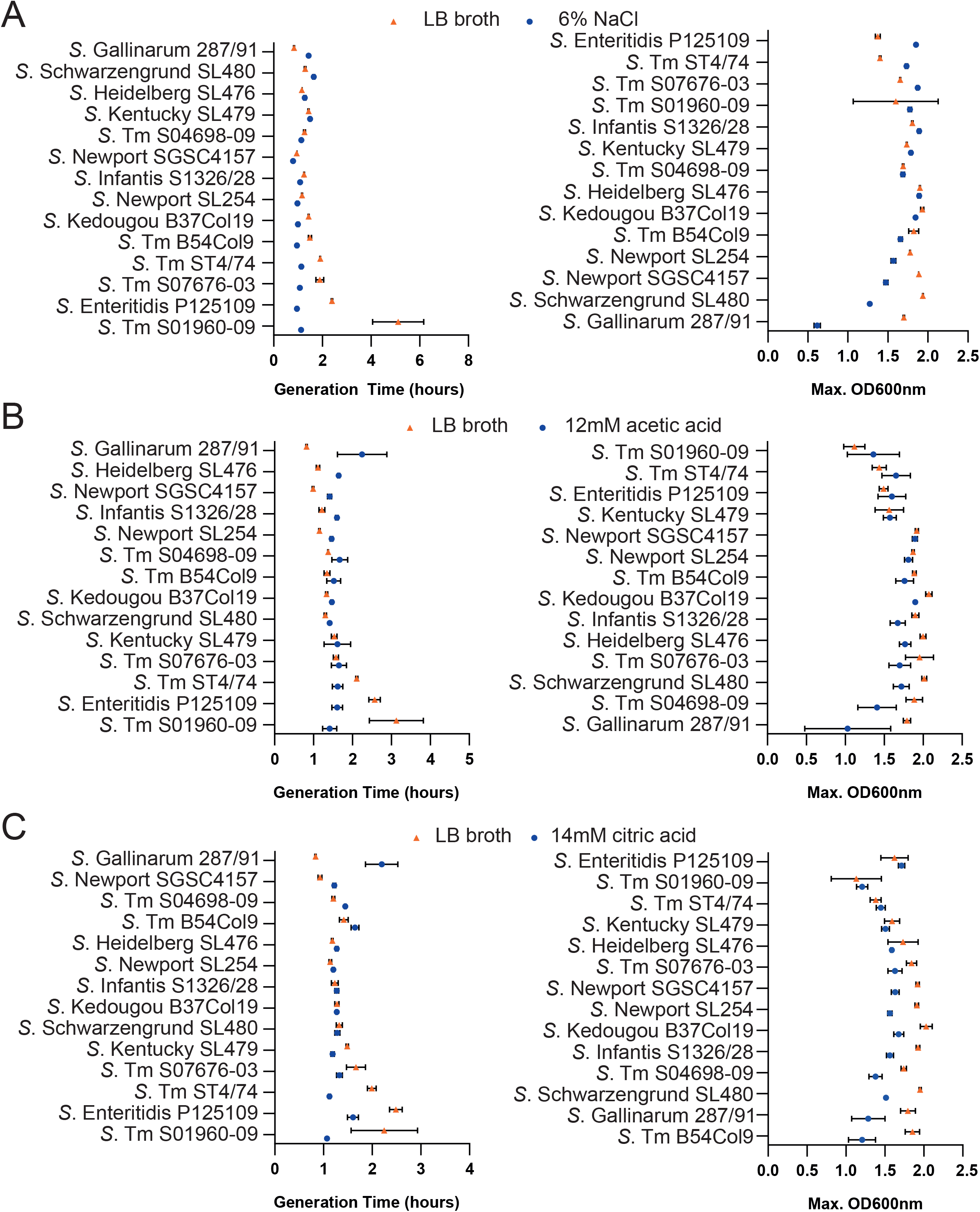
Minimum generation time and maximum optical density of *Salmonella enterica* strains in Luria-Bertani broth with or without supplementation with 6% NaCl, 12mM acetic acid or 14mM citric acid. Minimum generation time in log phase growth (left) and maximum optical density attained by cultures (right) of 14 *S. enterica* strains grown in LB broth (red triangle) or LB broth supplemented with (A) 6% NaCl, (B) 12mM acetic acid or (C) 14mM citric acid over a 24-hour period (blue circle). The mean of 3 biological replicates is plotted and standard error are shown.

The difference in AUC for each strain in 6% NaCl, 14mM citric acid and 12mM acetic acid was compared to the most resistant strain in each stress, which was *S*. Typhimurium strain S01960-05 for NaCl and acetic acid stress, and *S*. Enteritidis strain P125109 for citric acid (Figure 3). A two-way ANOVA revealed most strains had a significant difference in AUC compared to the most resistant strain in each stress (p < 0.05), except *S*. Typhimurium strain ST4/74 during citric and acetic acid stress, *S*. Enteritidis strain P125109 during NaCl and acetic acid stress and *S*. Typhimurium strain S01960-05 during citric acid stress (Figure 3).

## 4. Discussion

Consumer demand is rising for minimally processed food with fewer preservatives, and this needs to be delivered without compromising food safety. An improved understanding of variation in stress tolerance of diverse *Salmonella enterica* serotypes has the potential for improved risk assessment of processing techniques which satisfy consumer demand for minimally processed foods. Currently, strains of all *Salmonella* serovars are treated the same in risk assessments, however our data is consistent with the view that strains differ in their risk to food safety should they enter ready-to-eat food or if the food is insufficiently cooked.

In this study we investigated the variation in tolerance to food chain related stresses for 14 strains from nine different serovars of *Salmonella enterica* with diverse epidemiology and risk to food safety. Ten of the strains were amongst the top ten serovars most frequently isolated from human infection in 2019 in the UK (Enteritidis, Typhimurium, Newport, Infantis and Kentucky) (UKHSA, 2021). *S*. Enteritidis is generally associated with asymptomatic infection of poultry and eggs. Despite recent successful interventions that have resulted in a marked decrease in the incidence of *S*. Enteritidis in poultry in the UK, this serotype remains the most frequently isolated from human infection, presumably due to imported food products or travel. *S*. Infantis and *S*. Kentucky are emerging threats in the poultry industry, with MDR strains emerging in the last 10-20 years (EFSA and ECDC, 2017). *S*. Kentucky remains relatively rare in the UK, but outbreaks have occurred in the continental Europe associated with an MDR ST198 clone (Hawkey *et al.*, 2019). *S*. Newport is a minority serovar isolated from pigs and poultry yet is relatively frequently isolated from human infection in the UK (UKHSA, 2021). *S*. Kedougou is frequently isolated from UK poultry relative to other serovars, and although relatively rare in human infection, has caused several large outbreaks in multiple countries over the past 20 years (APHA, 2021). *S*. Heidelberg and *S*. Schwarzengrund have been linked to poultry and been associated with outbreaks including dried pet food (Behravesh *et al.*, 2010). *S*. Gallinarum is host restricted to poultry where it causes a severe disseminated infection called fowl typhoid. Occasional outbreaks in poultry flocks still occur in the high resource countries including the UK but is largely controlled by a test and cull programme. The epidemiology of *S*. Typhimurium is unusual in its widespread distribution in zoonotic reservoirs. Most human infections are from broad host range variants, currently typified by the monophasic *S*. Typhimurium ST34 clone that accounted for over half of all human *S*. Typhimurium infections in the past decade in the UK (EFSA, 2010; Moreno Switt *et al.*, 2009). Some variants exhibit varied degree of host adaptation. For example, *S*. Typhimurium U288 emerged in pigs in the UK around the same time as monophasic *S*. Typhimurium ST34 and despite accounting for similar proportions of pig isolates during this time, is relatively rarely associated with human infection. Significantly, *S*. Typhimurium U288 is more invasive in pigs and was present in the faeces in lower numbers than ST34 (Kirkwood *et al.*, 2021). *S*. Typhimurium DT56 circulate in populations of passerine (song) birds where they are thought to cause a severe disseminated infection (Lawson *et al.*, 2011). They can also occasionally be isolated from cattle and human infections (Mueller-Doblies *et al.*, 2018).

The strains investigated in this study were moderately diverse with average sequence divergence of nearly 2% for *S*. Schwarzengrund and *S*. Newport strain SGSC4157, but also included serovars *S*. Enteritidis and *S*. Gallinarum that differed by less than 0.3% and five strains of *S*. Typhimurium by less than 0.05% sequence divergence. However, genetic distance had little relationship to differences on observed tolerance to stresses tested. For example, the *S*. Gallinarum and *S*. Enteritidis strains had distinct tolerance to heat and cold storage and growth characteristics in 6% NaCl, 14mM citric acid and 12mM acetic acid despite their close relationship. Also, the *S*. Kentucky and *S*. Kedougou strains that were the next most closely related strains from different serovars also exhibited different levels of tolerance. Even *S*. Typhimurium strains had diverse tolerance to stresses, especially a *S*. Typhimurium U288 strain. Nonetheless, two strains of monophasic *S*. Typhimurium ST34 (S04698-09 and B54 Col9) differed by fewer than 20 SNPs in their entire genome and had similar tolerance to the stresses investigated. Together these observations suggest that differences in tolerance to stress can emerge relatively quickly but we did not find evidence of significant changes over very short periods of time during the ST34 epidemic clone that emerged in the past 20 years (Tassinari *et al.*, 2020).

Inappropriate food storage temperature is one of the most common causes of foodborne contamination (FAO and WHO, 2009). The recommended temperature for a domestic refrigerator is 2°C to 4°C to prevent microbial growth, however research into consumer refrigeration habits revealed that most household fridges exceed this temperature (FAO and WHO, 2009; Ovca *et al.*, 2021). None of the 14 strains tested in the current study increased significantly in viable count over a 5-week period indicating that the replication rate did not exceed the death rate in the food product at 4°C. This was expected since *Salmonella* typically grows between 7°C and 48°C, with an optimum temperature of 37°C, although some studies report growth at 2-4°C (Cox and Pavic, 2014; D’Aoust, 1991; Matches and Liston, 1968). *S*. Infantis strain S1326/28, *S*. Kedougou strain B37 Col19, *S*. Schwarzengrund strain SL480, *S*. Enteritidis strain P125109 and *S*. Typhimurium strains S07676-03, S01960-05, B54 Col9, S04689-09 all showed a greater tolerance to refrigerated temperatures. A similar study in egg yolks reported that *S*. Infantis counts at 5.5°C decreased by about 1-log over a 4-week period, although this may be due to the presence of antimicrobial properties of the food matrix (Lublin *et al.*, 2015). *Salmonella* is particularly problematic in low moisture foods such as dried fruit, peanut butter, and flour. Resistance to desiccation was variable amongst the 14 strains tested in the present study, consistent with a previous report of 37 strains of *Salmonella* in soybean meal (Norberto *et al.*, 2022). In the current study, both *S*. Typhimurium strain S01960-05 and *S*. Kentucky strain SL479 exhibited a lower tolerance. Tolerance to desiccation in U288 strains was also observed in a previous study whereby only 0.1% of cells of U288 strains remained viable after 24-hour desiccation (Kirkwood et al., 2021). In another study following *Salmonella* desiccation survival, *S*. Enteritidis had the highest tolerance (80% ± 9%) and *S*. Newport had the lowest survival rate (36% ± 3%) during 22-hour desiccation in a 96-well plate, which disagrees with the results from the present study where the mean log ratio survival for *S*. Enteritidis and *S.* Newport strain SL254 were similar, suggesting that tolerance to desiccation may be strain specific (Gruzdev *et al.*, 2011). Sodium chloride and organic acids are commonly used as a preservative to prevent microbial growth in food products, due to their cost effectiveness and ease of use (El Baaboua *et al.*, 2018), however our data suggests that *Salmonella* strains may be able to tolerate higher these chemicals to varying degrees and in some cases they may actually boost growth.

Notably, the strains that exhibit host adaptation tended to be the least tolerant to stress. *S*. Gallinarum strain 287/91 was the most sensitive to inactivation by heat and cold storage in food matrix, and *S*. Typhimurium strain U288 S01960-05 exhibited a relative low tolerance to desiccation compared to other serovars tested. *S*. Newport strain SGSC4157 also exhibited low tolerance to cold storage but was relatively stable during heat inactivation. A *S*. Typhimurium U288 strain was one of the least tolerant to heat inactivation and desiccation but was relatively stable during cold storage in food matrix. An exception to this trend was the relative tolerance to the stress conditions observed for a *S*. Typhimurium DT56 strain, that are adapted to passerine (song) birds. Adaptation to a single host or reduced host range has been reported to be associated with loss of function mutations in genes involved in multicellular behaviour, a mode of living thought to contribute to survival in the environment (MacKenzie *et al.*, 2017). For example, *S*. Typhimurium ST313 strains that are thought to be host adapted to an invasive lifestyle and have genomic signatures of host adaptation, have mutations affecting cellulose biosynthesis and catalase activity (MacKenzie *et al.*, 2019; Singletary *et al.*, 2016). A comparative analysis of *S.* Gallinarum strain 287/91 and *S.* Enteritidis strain P125109, revealed that *S.* Gallinarum harbours 309 hypothetically disrupted coding sequences (HDCSs) (Thomson *et al.*, 2008). These potential pseudogenes in 287/91 are responsible for loss of gene function in many metabolic pathways including, motility, metal/ drug resistance, amino acid catabolism and cellulose biosynthesis, and although not strictly related to the stresses included in this study, they could explain general sensitivity to environmental stress of *S.* Gallinarum. For example, strain 287/91 has a mutation in *bcsG* which may affect cellulose production, and therefore a lack of cellulose production also affects the cells’ ability to form biofilm, rendering it more sensitive to chemical and mechanical stress (Thomson *et al.*, 2008).

Growth of the host adapted strain of *S*. Gallinarum was also the most markedly retarded in the presence of 6% NaCl, 14mM citric acid and 12mM acetic acid. In contrast, growth of *S*. Typhimurium U288 was improved in the presence of these, although it should be noted that this strain grew poorly compared with all other strains tested in LB broth. The increased growth rate in the presence of 6% NaCl, 14mM citric acid and 12mM acetic acid was also observed for *S*. Enteritidis and *S*. Typhimurium strain ST4/74 and DT56. This is a similar finding to a previous study, in which an *S*. Enteritidis strain showed the smallest decrease in cell number during incubation with salt over a 3-hour period, and could resist the addition of salt concentrations as high as 8% (Wang *et al.*, 2020). The ability for *S*. Enteritidis to tolerate high salt concentrations poses a potential elevated threat in high-salt foods, such as soy sauce and seafood (Brown *et al.*, 2009).

The strains that exhibited the greatest tolerance to the stresses tested were two strains of monophasic *S*. Typhimurium ST34, that currently account for over half of all *S*. Typhimurium in people in the UK and over 10% of all *Salmonella* infections (UKHSA, 2021). These strains were among the most tolerant of cold storage and heat inactivation in food matrix and desiccation. Strains from other serovars commonly associated with human infection including *S*. Enteritidis, *S*. Newport and *S*. Infantis were all relatively tolerant of these stresses, except for *S*. Newport that was moderately but significantly less tolerant of cold storage compared to the most tolerant strain of *S*. Infantis. This relatively high level of tolerance may contribute to their prevalence in human infections. *S*. Kedougou was also notable for its relatively high tolerance to all stresses tested including 6% NaCl, 14mM citric acid and 12mM acetic acid having little impact on growth kinetics. Strains of *S*. Kedougou are also one of the most frequently isolated from poultry and poultry is known to be an important source of NTS infection, with over a quarter of all human Salmonellosis cases attributable to layer hens and broilers combined (EFSA, 2012). The virulence of strains of this serovar is demonstrated by its link to notable outbreaks and the recent past and therefore its relative scarcity may be due to other factors that have not been explored. For example, tissue tropism in poultry may affect the opportunity to enter the food chain, a factor that has been proposed to contribute to the limited the prevalence of *S*. Typhimurium U288 in human infection (Kirkwood et al., 2021).

## 5. Conclusions

The phenotypic variability in tolerance to a variety of food chain related stresses observed for strains of diverse *Salmonella enterica* serovars indicated that risk needs to be assessed on a strain-specific criteria. A greater understanding of the molecular basis of variation in tolerance to stress has the potential to identify molecular markers of risk that could be integrated into risk assessment of raw ingredients using information about processing and food-specific factors designed to inhibit or exclude *Salmonella.*

## Acknowledgements

RK, GT, LA and HK were supported by the BBSRC Institute Strategic Programme Microbes in the Food Chain BB/R012504/1 and its constituent projects BBS/E/F/000PR10348 and BBS/E/F/000PR10349. HP was supported by the UKRI Biotechnology and Biological Sciences Research Council Norwich Research Park Biosciences Doctoral Training Partnership as a National Productivity Investment Fund CASE Award (BB/R506126/1) with Nestlé.

---

**Supplementary Figure 1. Growth of *S*. Typhimurium strain ST4/74 in Luria-Burtani broth containing increasing concentrations of NaCl, citric acid and acetic acid.** Growth indicated by OD600nm is shown for ST4/74 cultures in LB broth containing increasing concentrations of (A) NaCl, (B) citric acid and (C) acetic acid. A representative experiment of two biological replicates is shown.

