## Supplementary figures and images for "Strain and serovar variants of *Salmonella enterica* exhibit diverse tolerance to food chain-related stress"

### Supplementary Table 1

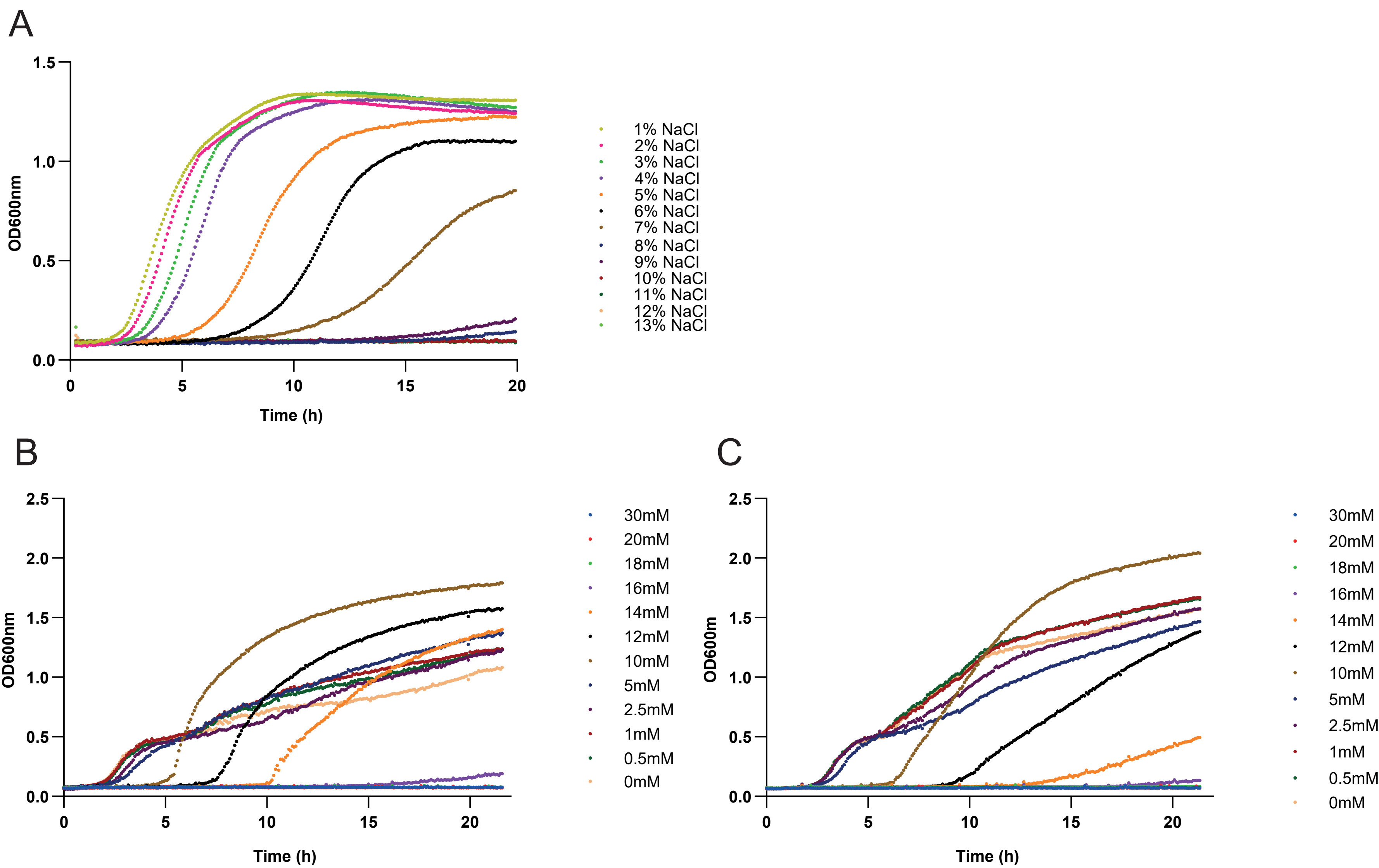
